# Description and comparison of PIMS-TS innate cell signature and immunophenotype with a cohort of healthy children, severe viral and bacterial infections and Kawasaki Disease

**DOI:** 10.1101/2021.03.29.437479

**Authors:** Alberto García-Salido, Inés Leoz-Gordillo, Anthony González Bravin, María Ángeles García-Teresa, Amelia Martínez de Azagra-Garde, María Isabel Iglesias-Bouzas, Marta Cabrero-Hernández, Gema De Lama Caro-Patón, José Luis Unzueta-Roch, Ana Castillo-Robleda, Manuel Ramirez-Orellana, Montserrat Nieto-Moro

**Author notes:** **Corresponding author**: Alberto García-Salido, M.D, PhD, Pediatric Intensive Care Unit. Hospital Infantil Universitario Niño Jesús. Avenida Menéndez Pelayo 65, Madrid, Spain. 34915035900. Email;. This study was partially funded by “Fundación Alonso”. The authors have disclosed that they do not have any potential conflicts of interest.

## Abstract

A new clinical syndrome associated to SARS-CoV-2 has been described in children. It has been named as Pediatric Inflammatory Multisystem Syndrome Temporally Associated with SARS-CoV-2 (PIMS-TS). This new disease is a main cause of hospital and pediatric intensive care unit (PICU). In this work we describe the innate cell signature and immunophenotype of children admitted to PICU because of PIMS-TS. Also, we compare it with healthy controls and children admitted to PICU because bacterial infection, viral infection and Kawasaki disease. We made a prospective-retrospective observational study in a tertiary pediatric hospital. Children admitted to PICU because of PIMS-TS from March 2020 to September 2020 were consecutively included. They were compare with previous cohorts from our center. A total of 247 children were included: 183 healthy controls, 25 viral infections, 20 bacterial infections, 6 Kawasaki disease and 13 PIMS-TS. PIMT-TS showed the lowest percentage of lymphocytes and monocytes with higher relative numbers of CD4+ (p =0,000). At the same time, we describe a differential expression of CD64, CD11a and CD11b. Monocytes and neutrophils in PIMS-TS showed higher levels of CD64 expression compared to all groups (p = 0,000). Also, proteins involved in leukocyte tissue migration, like CD11a and CD11b were highly expressed compare to other severe viral or bacterial infections (p = 0,000). In PIMS-TS this increased CD11a expression could be a sign of the activation and trafficking of these leukocytes. These findings are congruent with an inflammatory process and the trend of these cells to leave the bloodstream. In conclusion, we compare for the first time the innate cellular response of children with PIMS-TS with other severe forms of viral or bacterial infection and Kawasaki disease. Our findings define a differential cell innate signature. These data should be further studied and may facilitate the diagnosis and management of these patients.

---

A new type of affectation temporarily linked to the new coronavirus SARS-CoV-2 in childhood has been described. Europe, Great Britain and the United States have been the regions with the higher number of cases[1-3]. This new clinical syndrome has been named as Pediatric Inflammatory Multisystem Syndrome Temporally Associated with SARS-CoV-2 (PIMS-TS). It has clinical and analytical similarities to Kawasaki disease and suppose a main cause of hospital and pediatric intensive care unit (PICU) admission[4].

A majority of PIMS-TS cases are not related to active infections. An immune basis has been proposed for its establishment. In this point, there has been described the presence of immunity dysregulation or the release of autoantibodies. Related to that, immunoglobulin and corticoids seem to have a preeminent role. The clinical response is usually quick and with improvement in a short period of time[2,5].

Related to the probable role of leukocyte dysregulation the study of the cellular response in PIMS-TS could be of interest. Also, its comparison with healthy children and other causes of PICU admissions may help to describe and know the basis of their inflammation. Nowadays there are scarce data about this in children[6].

In this work we describe the innate cell signature and immunophenotype of children admitted to PICU because of PIMS-TS. Also, we compare it with healthy controls and children admitted to PICU because bacterial infection, viral infection and Kawasaki disease.

## Material and methods

Prospective-retrospective observational study conducted in a tertiary pediatric hospital after Ethics Committee for clinical research approval. Done prospectively in children admitted to PICU because of PIMS-TS from March 2020 to September 2020 were consecutively included. They were compare with previous cohorts from our center (see above). In all cases one peripheral blood sample was extracted after parents or legal guardians consent at PICU admission. A previously established intravenous line was always used. The volume obtained was 0.5 ml per sample and collected in sterile EDTA tube. The sample handling was based on item 59 of the Spanish law on Biomedical Research. Study was designed to not influence the treatment of the participating patients.

### Sample processing and analysis by flow cytometry

Samples were collected in sterile EDTA at room temperature or refrigerated at 4°C, used for CD45+ cells marking and analyzed by flow cytometry in a time period shorter than 24 hours. The antibody’s were all from Biolegend®: CD45 (clone HI30), CD4 (clone OKT4), CD8 (clone SK1), CD64 (clone 10.1), CD11a (clone TS2/4) and CD11b (clone M1/70). The surface expression were measured by BD FACS Canto II flow cytometer (Becton Dickinson, New York, USA). Cells viability were confirmed by 7-AAD staining. At least 10,000 events were recorded for each sample. The flow cytometer settings and samples were prepared according to the manufacturer’s instructions. Neutrophils, monocytes, and lymphocytes were identified on dot-plot profile and gated. The intensity of CD64 surface expression was measured as mean fluorescence intensity (MFI) in arbitrary units (monocytes as mCD64 and neutrophils as nCD64). The positive CD4, CD8, CD11a, CD11b and CD64 cells were expressed as percentage.

### Cohorts of analysis

Four cohorts of analysis were described:

1. Healthy patients: individuals under the age of 18 who came for programed surgery. They do not present previous diseases or infection. Sample extraction was performed prior to the intervention. Obtained in January 2018 to December 2019.
2. Viral and bacterial infections (VI and BI): individuals under 18 years of age admitted to PICU because of severe infection. Samples obtained at admission. Colonization was ruled out. Both groups are historical cohorts that have been described and published by our group before (see bibliography). *Definitions for a microbiologically confirmed case of bacterial infection included: 1) isolation of an organism by culture from blood, urine in a patient with clinical symptoms and signs of urinary tract infection or pyelonephritis, needle aspiration of abscess or empyema, stool sample of a patient with symptoms of gastroenteritis, pus sample from deep wound infection or detection of Streptococcus pneumoniae antigen from the pleural effusion sample of a patient with pneumonia.* *The diagnosis of a microbiologically confirmed viral infection required: 1) Detection of IgM antibodies or a fourfold increase in IgG antibodies in serum samples 2) viral antigen from nasopharyngeal aspirate or 3) viral nucleic acids by polymerase chain reaction of a nasopharyngeal aspirate, biological fluid or blood*.
3. Kawasaki disease (KD): children admitted to hospital or PICU which Kawasaki disease criteria. Period January 2018-december 2019. The flow cytometry was performed at hospital admission and prior to the initiation of immunomodulatory medication.
4. PIMS-TS: children admitted to PICU which met criteria defined for this condition (see bibliography). To confirm SARS-CoV-2 infection a nasal and pharyngeal swab using real-time reverse-transcriptase polymerase-chain-reaction (RT-PCR) was used. The presence of SARS-CoV-2 IgG antibodies was studied through ELISA.

### Statistical study

Statistical analysis was performed with the statistical program SPSS version 19.0 (IBM®). The quantitative values are expressed as mean and standard deviation. To compare quantitative variables between the bacterial and viral group a U Mann-Whitney test was used. Spearman’s rank correlation coefficient was bi-marginally calculated to measure the relationship between two continuous variables. A receiver operating characteristic (ROC) analysis with area under curve (AUC), sensitivity and specificity and cut-off values was performed for mCD64 and nCD64 to define its diagnostic accuracy for PIMS-TS. The cutt-off values were calculated by Youden index. Findings of two-tailed p < 0.05 were considered statistically significant.

## Results

A total of 247 children were included: 183 healthy controls, 25 viral infections, 20 bacterial infections, 6 Kawasaki disease and 13 PIMS-TS. In the PIMS-TS group 5 children were RT-PCR positive, 7 were Ig G positive and one RT-PCR and IgG positive at diagnosis. There were sex differences only in KD group. There were observed differences in age (Table 1). A negative correlation for lymphocytes with age was observed (r= - 0,240, p = 0,001). The CD64 expression in monocytes and neutrophils was not correlated with age or influenced by sex. The percentage of leukocytes and MFI of each type of leukocytes are described in table 1 and Figure 1. The significative differences between PIMS-TS and other groups are showed in Figure 1.

**Table 1.**
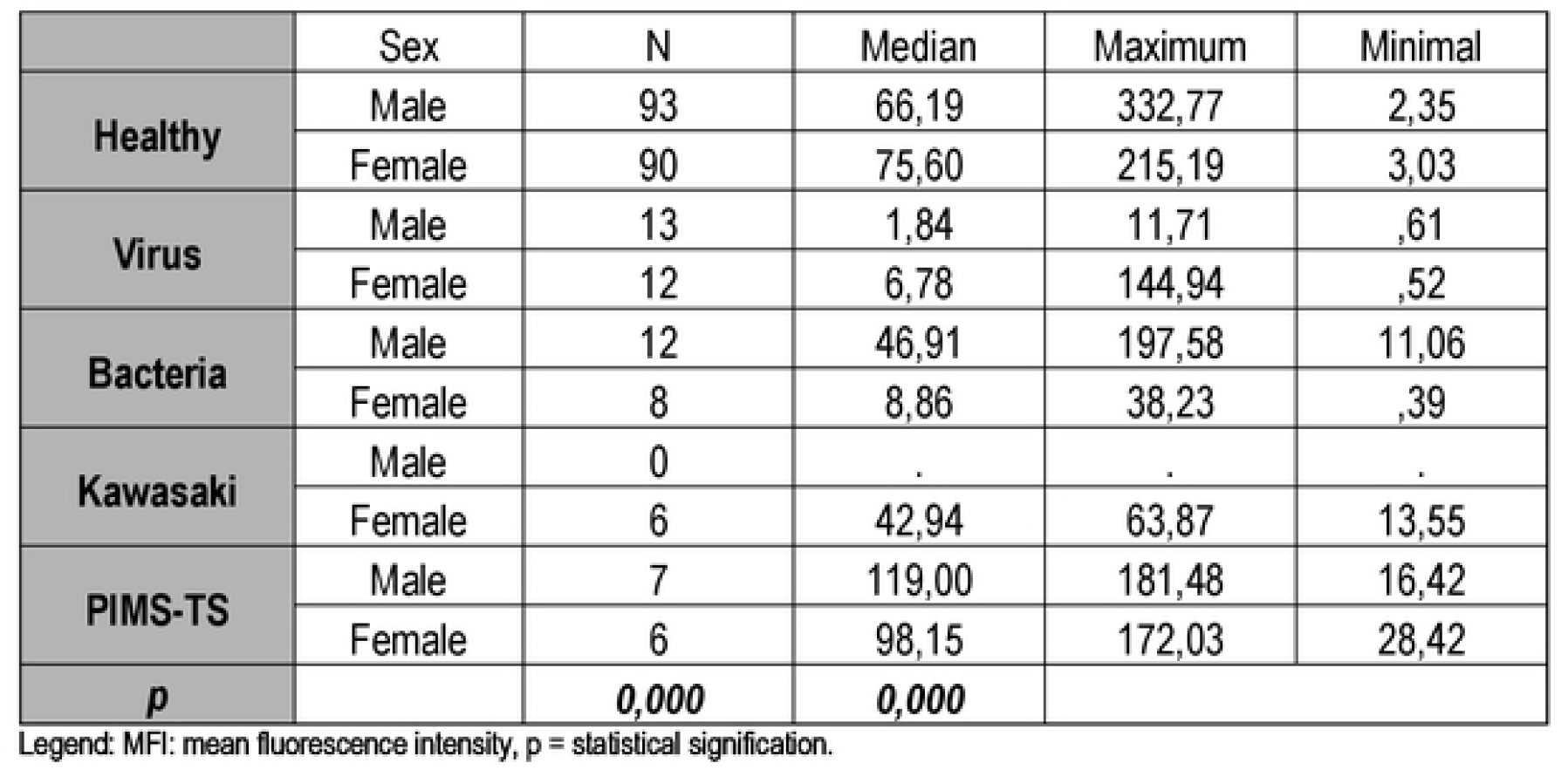
Age in months and sex of the cases included.

**Figure 1.**
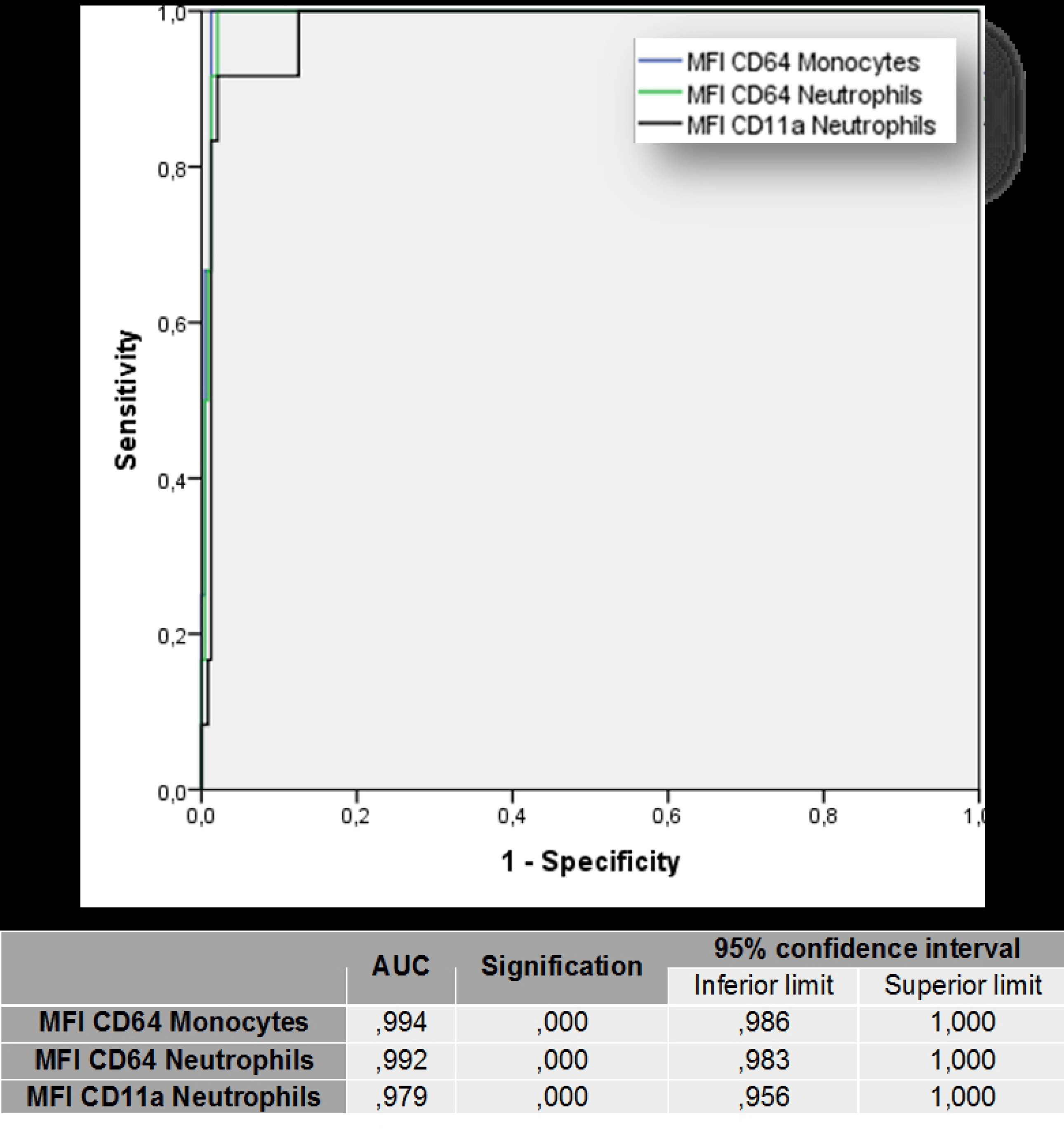
Leukocyte populations and immunophenotyping of patients included in the study. From left to right in each of the graphs: 183 healthy controls, 25 viral infections, 20 bacterial infections, 6 Kawasaki disease and 13 PIMS-TS. An arrow indicates those populations showing significant differences with PIMS-TS patients. **A**. Percentage of leukocyte populations. **B**. The first row compares the percentage of neutrophils positive for CD64, CD11a and CD11b. The bottom row compares the mean fluorescence intensity for CD64 in monocytes and neutrophils.

### CD64 and CD11a neutrophils expression utility as a PIMS-TS biomarker

To study the usefulness of CD64 surface expression as a tool to predict PIMT-TS we evaluated the receiver operating characteristics curve. As seen in Figure 2 the mCD64, nCD64 and nCD11a area under the curve (AUC) that were near to 1. For the mCD64 it was 0,994 (p=0,000; with a cut point of 29098 MFI and 94,1% specificity and 100% sensitivity). For the nCD64 it was 0,992 (p=0,000; with a cut point of 8753 MFI and 90% specificity and 100% sensitivity). The nCD11a AUC was 0,992 (p=0,000; with a cut point of 5203 MFI and 91,7% specificity and 94,1% sensitivity).

**Figure 2.**
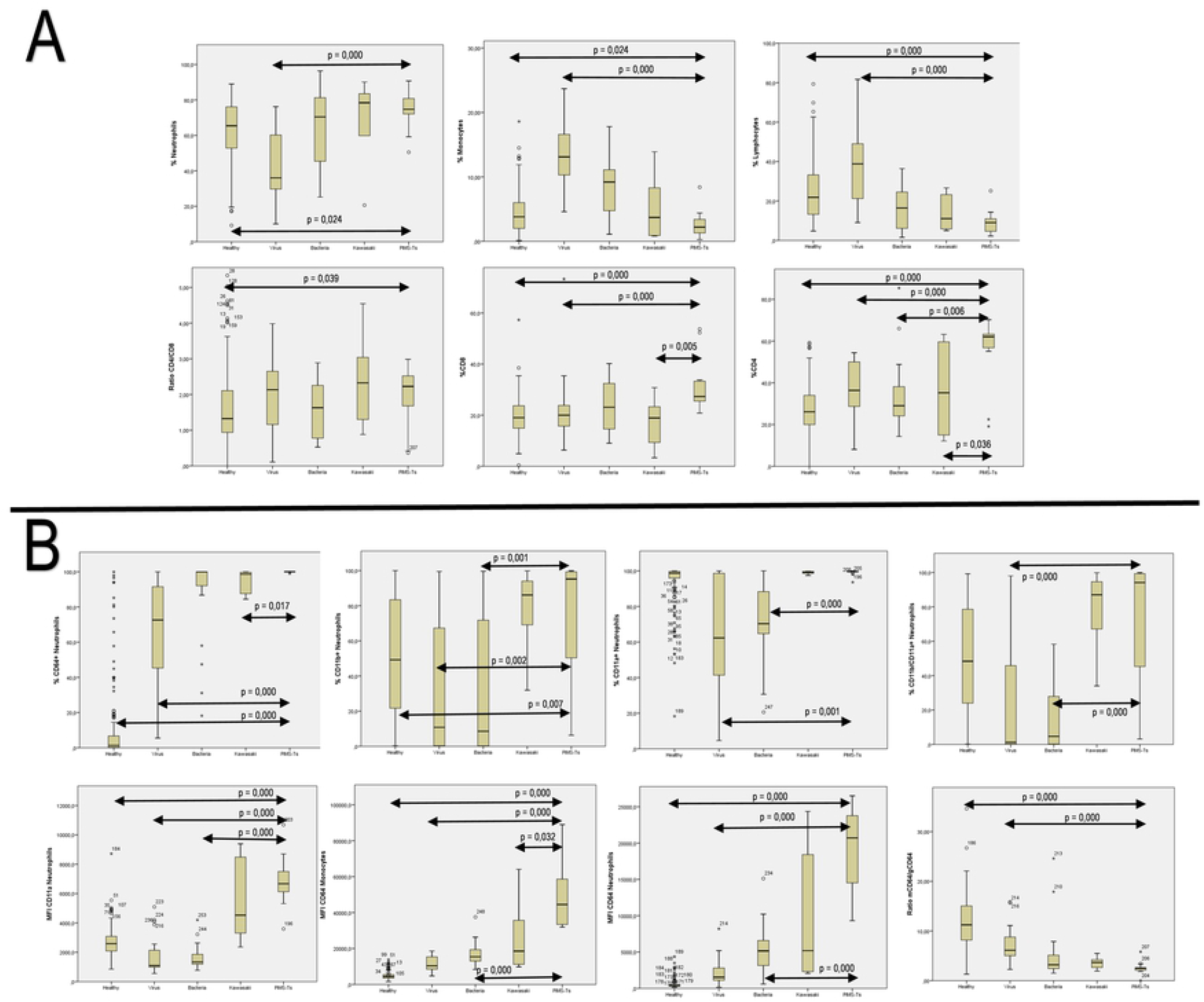
The ROC curve for CD64 on monocytes, CD64 on neutrophils and CD11a on neutrophils is shown. The values of the area under the curve with statistical significance and confidence interval are indicated.

## Discussion

This paper compares for the first time the innate cellular signature and immunophenotyping of severe PIMS-TS cases with a large cohort of healthy control, other severe infections diseases and KD. In peripheral blood samples, we observed differences in almost all leukocyte populations. The most visible of these differences affect lymphocytes and monocytes, which have the lowest values in PIMS-TS. At the same time, we describe a differential expression of CD64, CD11a and CD11b. These leukocyte surface proteins were exceptionally high in PIMS-TS in monocytes and neutrophils.

As said, the cases included showed a different distribution of leukocyte populations. This distribution of leukocyte populations is strikingly different in lymphocytes (Table 2 and Figure 1). PIMT-TS showed the lowest percentage of lymphocytes with higher relative numbers of CD4+. This lymphopenia has already been described in the adult population and severe critical children because of SARS-CoV-2 infections or PIMS-TS[2,7]. Compare to other severe viral infections, we observed that there was also a low percentage of monocytes and neutrophils (Figure 1). It is known that PIMS-TS is usually not associated with active SARS-CoV-2 infection. An immune dysfunction hypothesis has been proposed and explored in this children disease. Monocytes, neutrophils and lymphocytes are critical cells in viral first response. Their migration and accumulation in infected tissues added to the SARS-CoV-2 capacity to dysregulate this response may cause this low cell count in peripheral blood[6]. Our group has previously described an increase in leukocyte CD18/CD11a complex (LFA-1) expression in two short series of PIMS-TS[8,9]. It is congruent with a state of increased cellular predisposition to leave the bloodstream. We could not analyze the LFA-1 expression in this paper because it was not available in all groups.

**Table 2.**
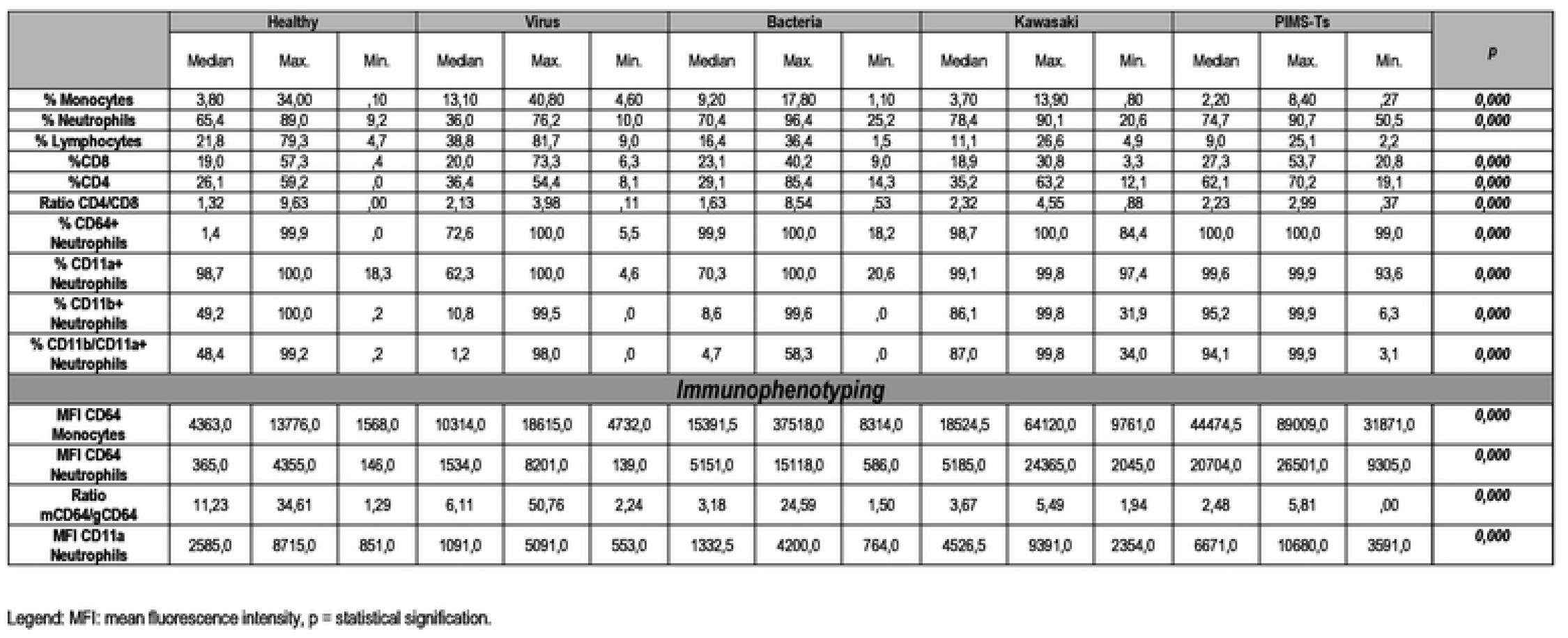
Comparison of leukocyte percentages and C064, C011a and C011b expression. The mean fluorescence intensity for CD64 and C011 a on neutrophils and monocytes is also described and compared.

Concerning immunophenotyping, we should highlight the findings observed about the expression of CD64, CD11a and CD11b. About CD64 expression, the PIMS-TS showed higher levels compared to all groups. The lowest CD64 value was observed in viral infections. The CD64 indirectly reflect cytokine expression and activation. As shown in Figure 2 the CD64 levels are higher in PIMS-TS than in severe bacterial infections and KD. This CD64 expression affects monocytes and neutrophils and may inform about the hyperinflammatory status in PIMS-TS[7,8]. Compared to KD we observed differences in monocytes but not in neutrophils. The study of CD64 and CD11a expression in these leukocytes could help in the differential diagnosis of PIMS-TS as is explained in Figure 2.

To study the proteins involved in leukocyte tissue migration, we examined the percentage of CD11a and CD11b positive cells. In our series, we observed that both proteins were higher in neutrophils and monocytes than in viral or bacterial infections (Figure 1). Also, they were higher than KD but without differences. Related to CD11a, the MFI was also higher in neutrophils of PIMS-TS cases. In adults, the inflammation in the basis of SARS-CoV-2 showed a predominant presence of macrophages and neutrophils in the affected territory. In PIMS-TS this increased CD11a expression could be a sign of the activation and trafficking of these leukocytes. These findings, added to the previously commented, are congruent with an inflammatory process and the trend of these cells to leave the bloodstream. This supports the inflammatory clinical findings but also confirm the utility of anti-inflammatory drugs as a cornerstone in the management of these children[10-13].

This work has limitations. We observed age differences between the groups analyzed (Table 1). The distribution of leukocyte populations is influenced by this. Despite this, we observed internal homogeneity between groups, which strengthens the observations. Also, the expression of CD64, CD11a and CD11b did not show correlations with age or differences by sex. Finally, the possible relationship between the observed data and the PIMS-TS clinical courses were not studied. This was not the aim of this work and should be considered in future works.

In conclusion, we compare for the first time the innate cellular response of children with PIMS-TS with other severe forms of viral or bacterial infection. Also, we describe differences with children with KD. Our findings define a differential cell innate signature with consistent data related to the possible presence of inflammation and immune dysregulation. These data should be further studied and may facilitate the diagnosis and management of these patients.

